## Supplemental Data for "A *Drosophila Su(H)* Model of Adams-Oliver Syndrome Reveals Notch Cofactor Titration as a Mechanism Underlying Developmental Defects"

**Supplemental Table S1**: Calorimetric Binding Data for Native and Variant RBPJ/Su(H) Proteins

| Ligand | Macromolecule | ΔG° (kcal/mol) | ΔH° (kcal/mol) | ΔS (kcal/mol(K)) | TΔS° (kcal/mol) | K ($M^{-1}$) | $K_D$ (nM) |
|---|---|---|---|---|---|---|---|
| DNA | Su(H) WT | -8.7 ± 0.1 | 8.5 ± 0.2 | 60.8 ± 0.3 | 17.2 ± 0.1 | 5.4 ± 0.8 x $10^6$ | 188.5 ± 27.4 |
| | Su(H) E137V | -7.9 ± 0.1 | 10.1 ± 0.5 | 63.4 ± 1.7 | 18.0 ± 0.5 | 1.2 ± 0.1 x $10^6$ | 841.6 ± 98.2 |
| | Su(H) K132M | -8.0 ± 0.1 | 11.2 ± 0.4 | 67.7 ± 1.2 | 19.2 ± 0.3 | 1.5 ± 0.1 x $10^6$ | 689.9 ± 48.7 |
| DNA | RBPJ WT | -9.2 ± 0.1 | 7.8 ± 0.1 | 60.1 ± 0.0 | 17.0 ± 0.0 | 1.4 ± 0.2 x $10^7$ | 74.0 ± 10.8 |
| | RBPJ E89G | -8.2 ± 0.1 | 12.0 ± 0.8 | 71.5 ± 2.5 | 20.2 ± 0.7 | 2.4 ± 0.5 x $10^6$ | 440.5 ± 86.8 |
| | RBPJ K195E | -7.7 ± 0.1 | 5.7 ± 0.9 | 47.3 ± 2.9 | 13.4 ± 0.8 | 9.0 ± 2.0 x $10^5$ | 1163.7 ± 258.1 |
| dNotch RAM | Su(H) WT | -9.2 ± 0.0 | -13.2 ± 0.1 | -13.5 ± 0.3 | -4.0 ± 0.1 | 5.4 ± 0.2 x $10^6$ | 186.7 ± 8.3 |
| | Su(H) E137V | -9.3 ± 0.2 | -13.3 ± 0.7 | -13.6 ± 2.8 | -4.1 ± 0.8 | 6.5 ± 1.7 x $10^6$ | 166.3 ± 47.4 |
| | Su(H) K132M | -9.6 ± 0.3 | -16.2 ± 3.5 | -22.1 ± 12.4 | -6.6 ± 3.7 | 1.2 ± 0.6 x $10^7$ | 106.7 ± 61.3 |
| mNotch1 RAM | RBPJ WT | -10.5 ± 0.0 | -12.5 ± 0.2 | -6.9 ± 0.6 | -2.1 ± 0.2 | 4.8 ± 0.3 x $10^7$ | 20.8 ± 1.2 |
| | RBPJ E89G | -10.6 ± 0.2 | -12.1 ± 0.1 | -5.2 ± 0.9 | -1.5 ± 0.3 | 5.6 ± 1.6 x $10^7$ | 19.9 ± 7.0 |
| | RBPJ K195E | -10.5 ± 0.2 | -12.2 ± 0.3 | -6.0 ± 1.4 | -1.8 ± 0.4 | 5.1 ± 1.8 x $10^7$ | 22.4 ± 7.8 |
| Hairless | Su(H) WT | -11.7 ± 0.1 | -16.2 ± 1.3 | -15.1 ± 4.7 | -4.5 ± 1.4 | 3.6 ± 0.4 x $10^8$ | 2.8 ± 0.3 |
| | Su(H) E137V | -12.1 ± 0.5 | -16.1 ± 1.4 | -13.5 ± 6.3 | -4.0 ± 1.9 | 1.1 ± 0.9 x $10^9$ | 1.7 ± 1.0 |
| | Su(H) K132M | -11.8 ± 0.8 | -16.5 ± 0.2 | -15.8 ± 2.7 | -4.72 ± 0.8 | 7.9 ± 5.9 x $10^8$ | 5.1 ± 5.9 |
| SHARP | RBPJ WT | -12.0 ± 0.9 | -11.7 ± 0.5 | 1.1 ± 1.3 | 0.3 ± 0.4 | 1.5 ± 1.7 x $10^9$ | 3.5 ± 3.3 |
| | RBPJ E89G | -11. ± 0.4 | -10.5 ± 1.4 | 3.0 ± 5.8 | 0.9 ± 1.7 | 2.9 ± 1.8 x $10^8$ | 5.2 ± 3.0 |
| | RBPJ K195E | -10.5 ± 0.3 | -13.3 ± 0.9 | -9.3 ± 3.8 | -2.8 ± 1.1 | 5.8 ± 2.4 x $10^7$ | 21.1 ± 10.1 |

**Supplemental Figure 1.** Differential scanning fluorimetry of purified WT and variant Su(H) (top) and RBPJ (bottom) proteins. The mean melting temperature (Tm) and standard error were calculated from triplicate experiments.

**Supplemental Figure 2.** EMSA from quick coupled *in vitro* transcription/translation system (TNT). Full length Flag-Rbpj WT (lane 1 and 2) binds to DNA in contrast to Flag-Rbpj K195E mutant (lane 5 and 6) and Flag-Rbpj E89G mutant (lane 3 and 4), which interact weakly with DNA. Protein-DNA interaction of Flag-Rbpj WT is shown by complex A (single occupancy, lane 1 and 2), complex A' (double occupancy, lane 1 and 2) and complex B and B' with the addition of anti-Flag showing supershifting (lane 2). Bottom panel: Western blot to show similar expression levels of WT and variant Rbpj constructs.

**Supplemental Figure 3.** Cycloheximide chase analysis of Myc-tagged RBPJ and RBPJ variant proteins. MK4 cells were transfected with Myc-tagged RBPJ constructs (human wild-type RBPJ, E63G, and K169E) followed by cycloheximide treatment (0.5ug/ml) for 24, 48, and 72 hours. Ectopic expression of Myc-tagged RBPJ constructs was detected by western blot analysis with anti-Myc (9B11) and anti-RBPJ antibodies. The signal intensities of corresponding protein bands were quantified by ImageJ software and expressed as ratios relative to those at day 1.

**Supplemental Figure 4.** Mammalian two-hybrid assay to analyze WT and Rbpj AOS variants binding to SHARP in cells. HeLa cells were cotransfected with the indicated Gal4-SHARP (300ng) and Rbpj-VP16 (50ng) constructs together with the pFR-Luc reporter (500ng), which contains five Gal4 DNA binding sites upstream of the luciferase gene. Relative luciferase activity was determined after cotransfection of the pFR-Luc reporter construct alone. WT, E89G, and K195E activate the reporter similarly, suggesting that the AOS variants bind SHARP similar to WT Rbpj, whereas the Rbpj double-mutant F261A/L388A, which is compromised for SHARP binding, does not activate the reporter.

**A**

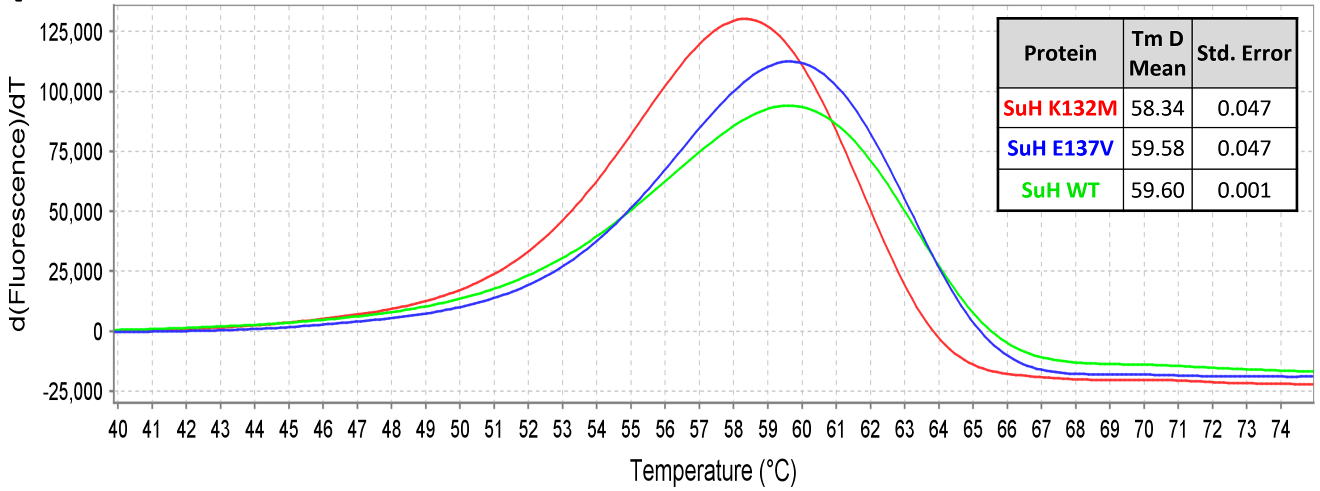


| Protein | Tm D Mean | Std. Error |
|---|---|---|
| SuH K132M | 58.34 | 0.047 |
| SuH E137V | 59.58 | 0.047 |
| SuH WT | 59.60 | 0.001 |

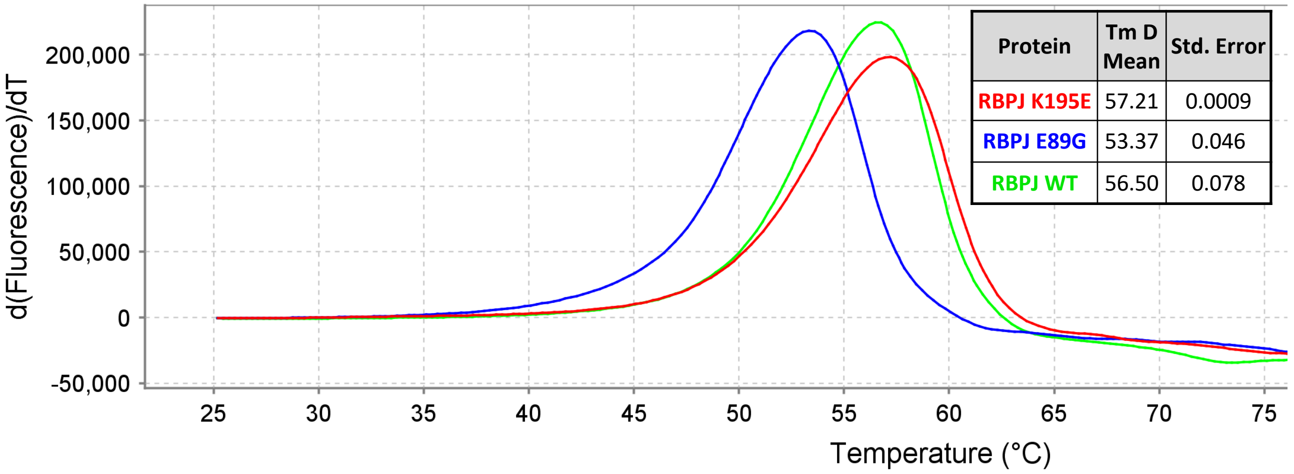


| Protein | Tm D Mean | Std. Error |
|---|---|---|
| RBPJ K195E | 57.21 | 0.0009 |
| RBPJ E89G | 53.37 | 0.046 |
| RBPJ WT | 56.50 | 0.078 |

**Fig S1**

**A**

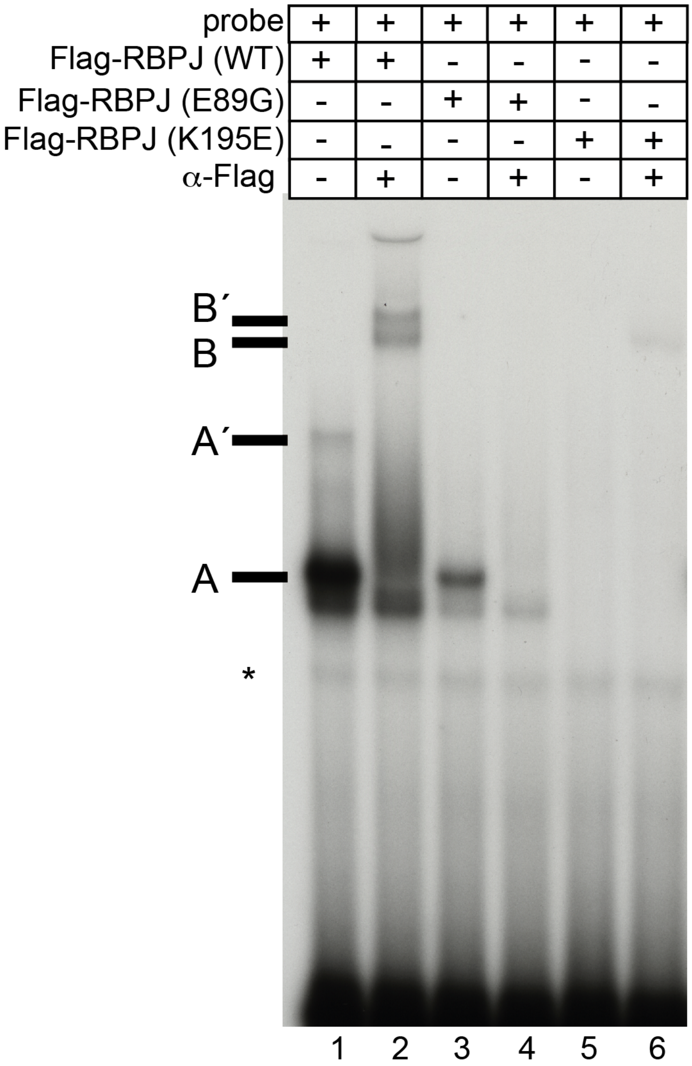
probe + + + + + +
Flag-RBPJ (WT) + + - - - -
Flag-RBPJ (E89G) - - + + - -
Flag-RBPJ (K195E) - - - - + +
α-Flag - + - + - +
B´
B
A´
A
*
1 2 3 4 5 6

**B**

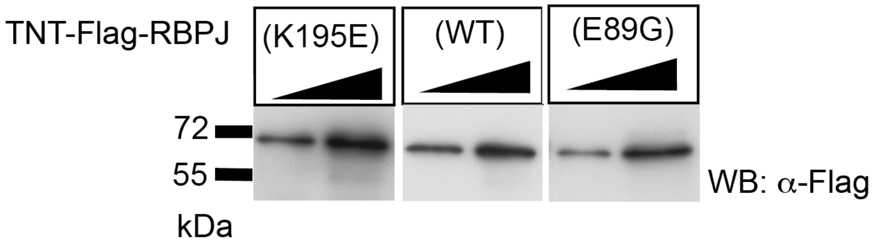
TNT-Flag-RBPJ
(K195E)
(WT)
(E89G)
72
55
kDa
WB: α-Flag

**Fig S2**

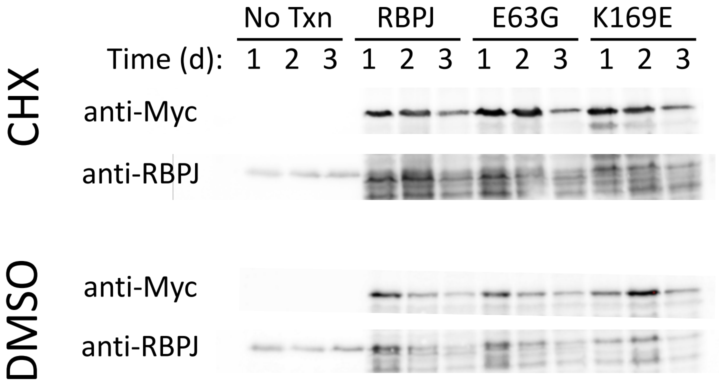
No Txn
RBPJ
E63G
K169E
Time (d): 1 2 3 1 2 3 1 2 3 1 2 3
CHX
anti-Myc
anti-RBPJ
DMSO
anti-Myc
anti-RBPJ

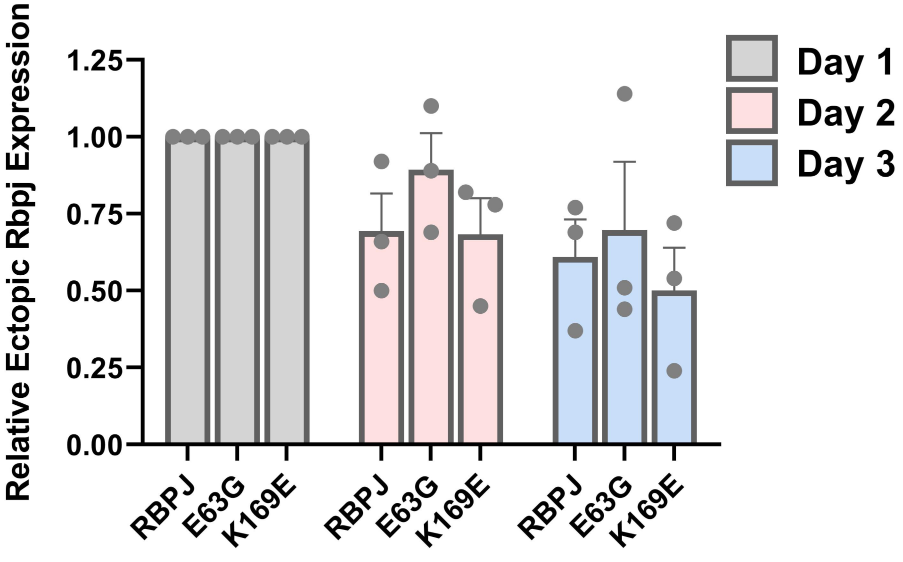
Relative Ectopic Rbpj Expression
1.25
1.00
0.75
0.50
0.25
0.00
Day 1
Day 2
Day 3
RBPJ
E63G
K169E
RBPJ
E63G
K169E
RBPJ
E63G
K169E

**Fig S3**

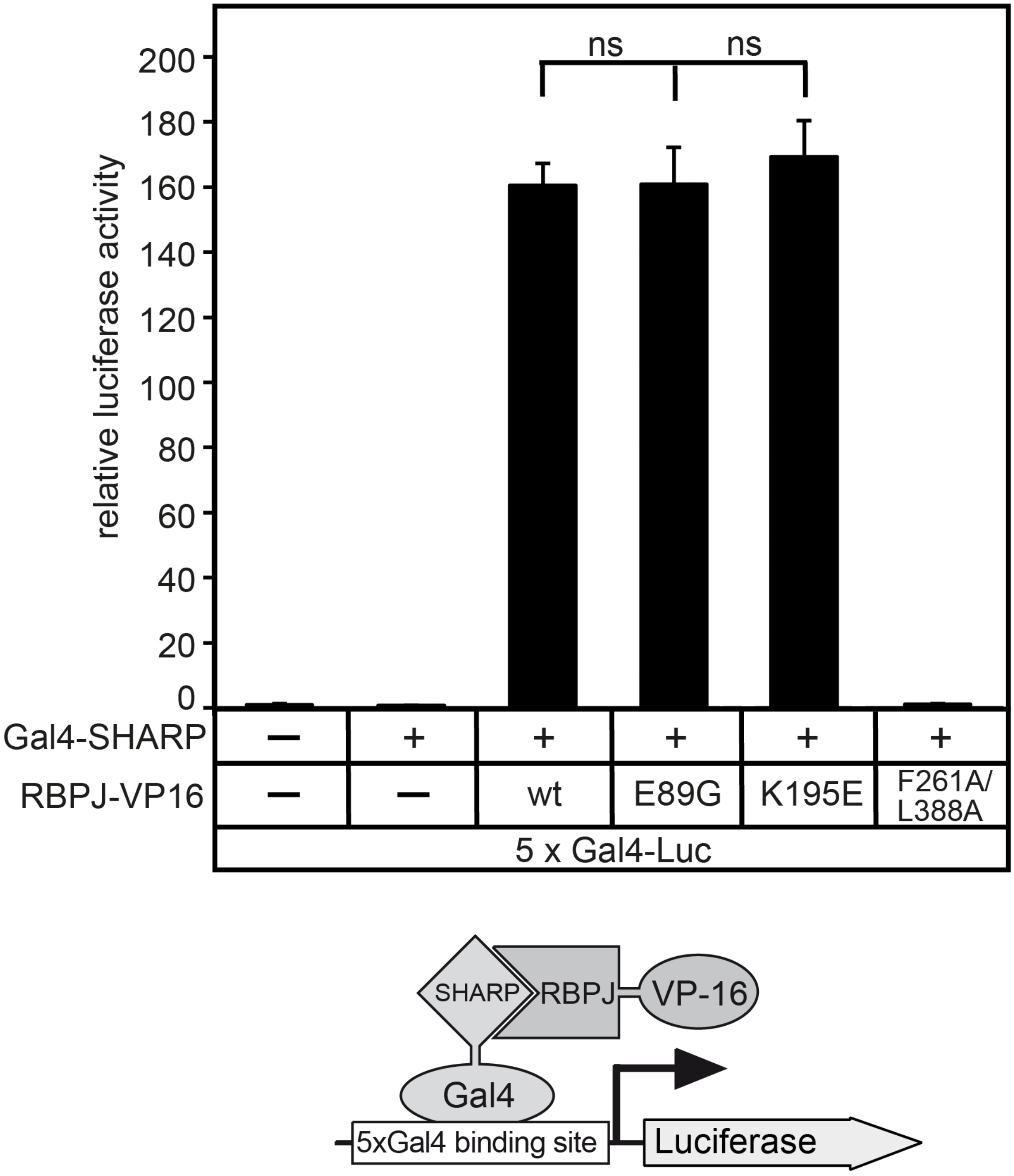
relative luciferase activity
200
180
160
140
120
100
80
60
40
20
0
ns
ns
Gal4-SHARP
RBPJ-VP16
wt
E89G
K195E
F261A/
L388A
5 x Gal4-Luc
SHARP
RBPJ
VP-16
Gal4
5xGal4 binding site
Luciferase

**Fig S4**
